# Novel murine closed-loop auditory stimulation paradigm elicits macrostructural sleep benefits in neurodegeneration

**DOI:** 10.1101/2024.02.12.579922

**Authors:** Inês Dias, Sedef Kollarik, Michelle Siegel, Christian R. Baumann, Carlos G. Moreira, Daniela Noain

**Affiliations:** Department of Neurology, University Hospital Zurich (USZ), Switzerland; Department of Health Sciences and Technology (D-HEST), ETH Zurich, Switzerland; Neuroscience Center Zurich (ZNZ), Switzerland; Center of Competence Sleep and Health, University of Zurich (UZH), Switzerland

**Keywords:** Mouse closed-loop auditory stimulation, non-rapid eye movement sleep, neurodegeneration, Alzheimer’s disease, Parkinson’s disease

## Abstract

Boosting slow-wave activity (SWA) by modulating slow-waves through closed-loop auditory stimulation (CLAS) might provide a powerful nonpharmacological tool to investigate the link between sleep and neurodegeneration. Nevertheless, CLAS in this context was not yet explored. Here, we established mouse CLAS (mCLAS)-mediated SWA enhancement and explored its effects onto sleep deficits in neurodegeneration, by targeting the up-phase of slow-waves in mouse models of Alzheimer’s (AD, Tg2576) and Parkinson’s disease (PD, M83). We found that tracking a 2Hz component of slow-waves leads to highest precision of NREM sleep detection in mice, and that its combination with a 30° up-phase-target produces a significant SWA 15-30% increase from baseline in WT_AD_ and TG_AD_ mice versus a MOCK group. Conversely, combining 2Hz with a 40° phase target yields a significant increase ranging 30-35% in WT_PD_ and TG_PD_ mice. Interestingly, these phase-target-triggered SWA increases are not genotype dependent but strain specific. Sleep alterations that may contribute to disease progression and burden were described in AD and PD lines. Notably, pathological sleep traits where rescued by mCLAS, which elicited a 14% decrease of pathologically heightened NREM sleep fragmentation in TG_AD_ mice, accompanied by a steep decrease in microarousal events during both light and dark periods. Overall, our results indicate that model-tailored phase-targeting is key to modulate SWA through mCLAS, prompting the acute alleviation of key neurodegeneration-associated sleep phenotypes and potentiating sleep regulation and consolidation. Further experiments assessing the long-term effect of mCLAS in neurodegeneration may majorly impact the establishment of sleep-based therapies.

## Introduction

Auditory stimulation has recently emerged as a promising approach to modulate sleep slow-waves in a precise manner (Wilckens et al., 2018). When auditory stimuli are timed or phase-locked to spontaneous positive (up-phase) waves during slow-wave sleep (SWS), an increase in slow-waves and spindles is observed, which correlates with improved memory recall in healthy subjects, patients with amnesic mild cognitive impairment, and children with attention deficit hyperactivity disorder (Ngo et al., 2013; Papalambros et al., 2017, 2019; Prehn-Kristensen et al., 2020). Conversely, targeting the down-phase of slow-waves impairs neuroplasticity and learning in healthy subjects (Fattinger et al., 2017; Moreira et al., 2021). Even though it has been deemed effective as stand-alone acute intervention in brain injury (Moreira et al., 2022), it remains elusive whether CLAS would be attainable and beneficial in the context of chronic brain disease, such as neurodegeneration.

Sleep-wake disturbances are a common feature of the asymptomatic preclinical stages of Alzheimer’s (AD) and Parkinson’s disease (PD), suggesting sleep may play a key role in the onset and progression of neurodegenerative diseases (Holth et al., 2019; J. E. Kang et al., 2009; Lucey, 2020; Minakawa et al., 2017; Owen & Veasey, 2020; Qiu et al., 2016). Thus, a bidirectional relationship between sleep and neurodegeneration is currently acknowledged, in which disrupted sleep - particularly delta-rich SWS - results from the degeneration of sleep regulatory brain regions and, inversely, neurodegeneration progression is worsened by SWS disruption (Ju et al., 2014; Mapelli, 2015; Schreiner et al., 2019; Wang & Holtzman, 2019). Critically, pharmacological sleep induction protocols were associated with decreased amyloid-beta or alpha-synuclein load in AD and PD mouse models, respectively (J. Kang et al., 2009; Morawska et al., 2021). However, similar pharmacological approaches aiming at SWS increase in patients (Büchele et al., 2018; Mednick et al., 2013; Walsh, 2009), even though effective, present major limitations such as the high risk of tolerance, dependency, misuse and lack specificity (Cellini & Capuozzo, 2018; Lie et al., 2015). In this context, if successfully established at preclinical level, CLAS could become a tool of choice to enhance SWS in patient populations for it is regarded as a reliable, safe, and relatively noninvasive to apply in a home-based environment, which may be ideal for patients displaying cognitive or motor impairments (Ferster et al., 2019; Lustenberger et al., 2022).

Despite these remarkable positive prospects, the lack of animal models implementing CLAS in neurodegeneration, thus allowing the combination of phase-targeted sound-evoked neuromodulation in a chronic disease context, has delayed its optimization and translation into clinical settings. Therefore, establishing whether CLAS is able to successfully modulate slow-waves in mouse models of neurodegeneration is a timely and crucial endeavor, enabling critical further assessments on whether this technology may be beneficial in rescuing disturbed sleep phenotypes in early stages of neurodegeneration.

Here, we successfully established up-phase-targeted mouse CLAS (mCLAS) in two mouse models of neurodegeneration: 7-month old AD mice, Tg2576 line (Hsiao et al., 1996); and 7-month old PD mice, M83 line (Giasson et al., 2002). Our aims comprised developing a novel mCLAS methodology enabling mechanistic and therapeutic assessments of sleep-mediated neuroprotection in neurodegeneration, and assessing mCLAS’s short-term effects onto neurodegeneration-associated sleep disturbances.

## Methods

### Animals

We used the Tg2576 model of AD (Hsiao et al., 1996), overexpressing a mutant form of the human amyloid precursor protein (APP) gene, the APPK670/671L; and the M83 model of PD (Giasson et al., 2002), expressing the A53T-mutated human variant of α-synuclein gene under the direction of the mouse prion protein promoter. The Tg2576 line was bred in the B6;SJL-Tg(APPSWE)2576Kha strain (B6;SJL Mixed Background) whereas the M83 line was bred in the B6.Cg-2310039L15Rik^Tg(Prnp-SNCA*A53T)23Mkle^/J strain (C3H/HeJ x C57BL6/J hybrids). The study design included female and male TG mice of each model (TG_AD_ and TG_PD_) at a prodromal disease stage (6.5 months-of-age when implanted, plaque- or heavy inclusion-free) and their WT littermates (WT_AD_ and WT_PD_). Females amounted to 60% of the animals used (**Supplementary Table 1**). Mice were group-housed in ventilated cages with food and water *ad libitum*, under a 12/12h light/dark cycle. All experimental procedures were performed according to federal Swiss regulations under approval and supervision of the veterinary office of the Canton Zurich (license number ZH024/2021).

### Surgery

Mice were implanted as described before (Kollarik et al., 2022; Morawska et al., 2021). Briefly, we placed 1 stainless steel screw per hemisphere, each located 2 mm posterior to Bregma and 2 mm lateral from midline. The screws were connected to a pin header via copper wires for EEG recordings, and the structure fixed with dental cement. For EMG recordings, we inserted two gold wires bilaterally into the neck muscles (**Figure 1A**). All surgical procedures were performed under deep inhalation (with isoflurane, 4-4.5% for induction in anesthesia box and 2-2.5% for maintenance using a nose cone fitting) and local (lidocaine) anesthesia, with administration of anti-inflammatory (5 mg/kg, s.c., Metacam®) and analgesic drugs (0.1 mg/kg, s.c., Temgesic®). Postsurgical analgesia was administered for two days during both light and dark periods, with body weight and home cage activity daily monitored during the surgical week and twice weekly thereafter.

**Figure 1.**
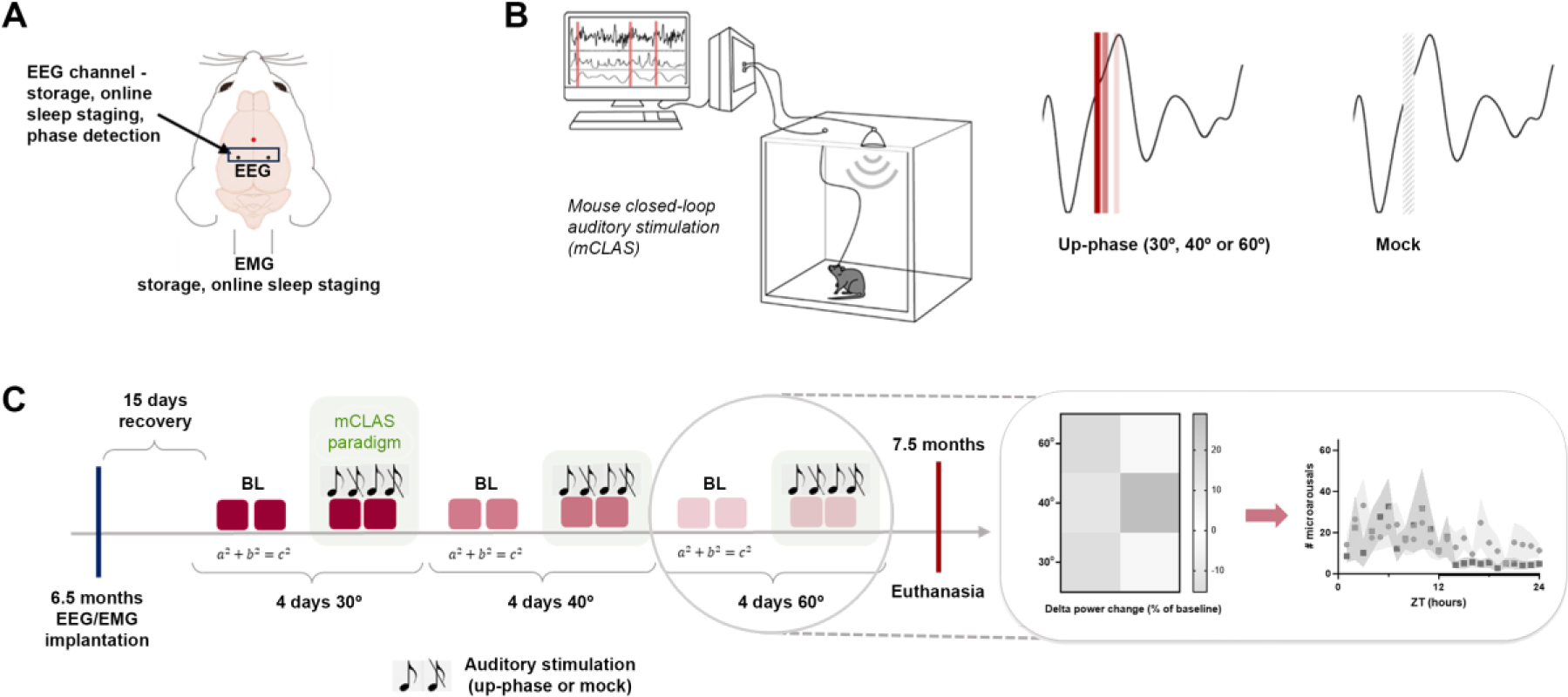
Experimental design. **A:** Dorsal skull schematic of electrode placement upon surgery. **B:** After recovery, animals are accommodated in individual chambers for either 30°, 40° or 60° up-phase stimulation (with sound delivery) or mock stimulation (sounds muted but events flagged). **C:** Experimental timeline depicting the sequence of events. During baseline (BL) days, threshold calculations were performed. Analysis was divided in two main parts: we first accessed technical parameters of mCLAS performance and its effect on delta power; secondly, we explored the short-term effects of our stimulation paradigm on disease-related sleep deficiencies.

### EEG/EMG recordings

Following a two-week post-surgical recovery, mice were individually housed in custom-built sound-insolated chambers, which simultaneously and continuously record EEG/EMG, monitor behavior (through an infrared recording camera in each chamber) and deliver phase-locked auditory stimuli to seven mice independently and in real-time (Moreira et al., 2021). We conducted intra-hemispheric tethered recordings (differential mode), with data acquired through a Multi-channel Neurophysiology System (TDT, USA). Left hemisphere EEG signal was referenced to the right hemisphere electrode, amounting for one EEG channel per mouse. We recorded all signals using SYNAPSE software (TDT), which were subsequently sampled at 610.35 Hz, amplified (PZ5 Neuro-Digitizer preamplifier, TDT) after applying an anti-aliasing low-pass filter (45% of sampling frequency), synchronously digitized (RZ2 BIOAMP processor, TDT), and finally stored locally (WS-8 workstation, TDT). We filtered real-time EEG between 0.1-36.0 Hz (2^nd^-order biquad filter), and EMG between 5.0-525.0 Hz (2^nd^-order biquad filter and 40 dB notch filter centered at 50 Hz). These signals served as input to real-time detection algorithms for stimuli delivery.

### mCLAS paradigm

We performed the experiments in batches of 5-7 mice, in a 12/12h light/dark cycle and with water and food available a*d libitum*. Upon connection to the tether, implanted mice (**Figure 1A**) were given 2 days to adapt to the chamber and cabling. In WT and TG MOCK groups, possible stimulation events were flagged but sound stimuli was not delivered throughout the procedure. In WT and TG stimulation groups, we adapted the paradigm developed previously for rats (Moreira et al., 2021). Briefly, auditory stimuli consisted of clicks of pink 1/f noise (15ms, 35 dB SPL, 2ms rising/falling slopes) delivered by inbuilt speakers on top of the stimulation chamber. We determined 15ms stimuli duration as ideal through pilot studies in WT mice, where 15, 20, 25 and 30ms pulses were tested (data not shown). Such finding is consistent with literature suggesting that mice slow-waves have a shorter cycle than those of rats and humans (Mölle et al., 2009; Panagiotou et al., 2017), meaning short durations are required for proportional stimuli overlapping.

For phase-targeted mCLAS, a nonlinear classifier defined four decision nodes comprising power in EEG (EEGpower), power in EMG (EMGpower), a frequency component and a phase-target. For EEGpower, we computed the ratio between rms of delta (0.5-4Hz) and high-beta (20–30Hz) bands on a 1s sliding window (using an algorithm written in RPvdsEx, Real-Time Processor Visual Design Studio, TDT) and compared it to an EEGthreshold extracted from 48h EEG/EMG non-stimulated baseline recordings automatically scored and fed to a custom-written MATLAB R2020b script (Moreira et al., 2021). In short, an estimate of EEGpower during NREM was established as +1.0 SD above the mean (84.1% percentile) of the EEGpower of all NREM epochs during baseline. For EMGpower, we computed rms EMG and compared it to a baseline EMGthreshold, confined to values –1.0 SD below the mean (15.9% percentile) of the rms EMG values during offline scored NREM. EEGthreshold and EMGthreshold marked the transition into consolidated NREM sleep in each mouse and were introduced in our custom-made SYNAPSE project. Regarding the frequency component node, a very-narrow bandpass filter tracked a user-defined frequency component from the EEG channel. For phase, we defined slow-wave’s 0° as the rising zero-crossing and 90° as the positive peak. At every identified positive zero-crossing on the filtered signal, the phase-detector resets to 0° and calculates a user-defined phase-target based on the number of elapsed samples since 0⁰. To deliver sound stimuli, validation of truth value of the four decision nodes need to be simultaneously met, with a trigger delay of ∼29.30ms. All triggers were flagged and saved for offline analysis. We used the recording cameras to control for stimuli-evoked arousals or reflexes.

In WT and TG stimulation groups, we predefined 60⁰ of the slow-wave as phase-target and tracked either the 1, 1.5, 2 or 2.5Hz component from each subject’s EEG channel. In rats, 1Hz/60⁰ parameter set was ideal (Moreira et al., 2021), but as SWA peaks around 2-2.5Hz in mice (Mölle et al., 2009; Panagiotou et al., 2017), we here tracked frequencies ≥1Hz. After assessing and defining the ideal frequency component to be tracked in mice, we tested three up-phase-targets (30⁰, 40⁰ and 60⁰) in all genotypes and strains (**Figure 1B**). Within each condition tested, we sequentially recorded 2 EEG/EMG baseline days and 2 EEG/EMG stimulation days (**Figure 1C**) in each subject, with parameters’ order randomized. Recordings were conducted between 1-2 weeks from subjects’ connection.

### EEG/EMG offline scoring and online staging validation

We scored all recorded files using SPINDLE for animal sleep data (Miladinović et al., 2019), which was previously validated in our lab for rats and mice (Kollarik et al., 2022; Moreira et al., 2021). SPINDLE retrieved vigilance states (wakefulness, NREM and REM sleep) with 4s epoch resolution from EDF files with EEG/EMG signals. Wakefulness was characterized by high or phasic EMG activity for more than 50% of the epoch duration and low amplitude but high frequency EEG. NREM sleep was hallmarked by the presence of slow oscillations, reduced EMG activity and increased EEG power in the delta frequency band (0.5 - 4 Hz). REM sleep was defined based on high theta power (6 - 9 Hz band) and low EMG activity.

To validate our online staging method, we compared it with SPINDLE offline scores. Briefly, we grouped 1s epochs in 4s blocks, which were categorized as NREM sleep when ≥50% of 1s values were scored as such. To assess staging performance, we calculated a set of diagnostic parameters (precision, specificity and sensitivity) per day between online and offline staging approaches. Precision was computed as the proportion of online labeled NREM sleep epochs confirmed offline as NREM among all online NREM labels. Specificity was calculated as the ratio between online epochs confirmed offline as non-NREM sleep and the total non-NREM epochs detected offline, assessing if the online staging can exclude all that it is non-NREM. Sensitivity was calculated as the ratio between online NREM sleep epochs confirmed offline as NREM and the total NREM epochs detected offline, measuring how strictly the algorithm identified sustained NREM sleep using the thresholds defined.

### EEG/EMG offline pre- and post-processing

#### Delta power

Following online staging validation, we processed EEG data through a custom-made MATLAB script. In brief, we first removed artifacts by detecting clipping events (15 adjacent raw EEG samples of approximately 60ms within 55-units of the amplifier maximum/minimum), followed by a 3-point moving average (to remove frequencies ≥80Hz) and a basic Fermi window function (to gradually attenuate the first and last 2s of each signal, preventing ringing artifacts) (Modarres et al., 2017). Next, we resampled the signal at 300Hz and filtered between 0.5–30Hz using low and high-pass zero-phased equiripple FIR filters (Parks–McClellan algorithm applied in both directions, filtfilt). We inspected the signal for regional artefacts (2h sliding window) not detected during automatic scoring or the above mentioned steps: within scored NREM sleep, brief portions of the signal (<10 sample-points at 300 Hz) > ± [8×interquartile range] were reconstructed by piecewise cubic spline interpolation from neighboring points. We extracted delta power through a spectral analysis of consecutive 4s epochs (FFT routine, Hamming window, 2s overlap, 0.25Hz resolution) and normalized the data indicating the percentage of each bin with reference to the total spectral power between 0.5-30Hz, per 24h recording. Delta power changes per day of stimulation were calculated as percentage of change from baseline recordings of each animal.

#### Total NREM sleep time, NREM sleep fragmentation index and microarousals

As a metric of sleep architecture, we determined total time spent in NREM sleep in minutes, and total NREM sleep time as percentage of change from baseline for each mouse. NREM sleep fragmentation index was expressed as the number of NREM sleep bouts/total number of NREM sleep epochs, with higher index values expressing higher fragmentation. A NREM sleep bout was defined as the duration of consecutive NREM sleep epochs. We also calculated NREM sleep fragmentation index as percentage of change from baseline for each animal. NREM sleep time and fragmentation were both calculated based on the EEG signal pre-processed as described in the previous section. To attest for sleep consolidation, we calculated microarousals, defined as high-amplitude power bursts in EMG signal during NREM sleep upon artifact removal. In brief, after applying a basic Fermi window function, we resampled the signal at 300Hz and filtered between 5–100Hz using low and high-pass zero-phased equiripple FIR filters (same fashion as previously described) with a bandstop centered at 50Hz. Next, we applied a moving RMS function to the signal, with values inspected for power bursts: within scored NREM sleep epochs, brief portions of the signal (<10 sample-points at 300 Hz) > ± [7×interquartile range] were considered microarousal events when falling within the outlier quartiles, for less than 2 seconds. Microarousal events were grouped by hour in ZT allowing for a visualization of their 24h dynamics during both light and dark periods (plotted for every 3-hour). We also calculated the cumulative number of microarousals per 24h, light and dark periods separately, for each animal.

#### Auditory trigger distribution

We analyzed trigger distribution as 1) the daily number of triggers in NREM sleep and other stages (wakefulness and REM sleep) and 2) the ratio between the number of triggers in NREM sleep and the total number of triggers. To assess the ability of the paradigm to successfully target the up-phases tested, we filtered EEG data between 0.5-2Hz (same fashion as previous digital filters), followed by a Hilbert transform to define instantaneous phase. We averaged the instantaneous phase for all triggers delivered in NREM sleep and plotted the results in histograms of 10° bins. Counts were normalized to the total number of triggers in each animal.

### Statistical analysis

We conducted statistical analysis in IBM SPSS Statistics 28 (IBM, USA), with all data plotted in GraphPad Prism 10.0.2 (GraphPad Software, LLC). We inspected the data for outliers through boxplots, with outliers considered as values 1.5 box-lengths from the edge of the box.

The distribution of each variable and its residuals was assessed for normality through skewness and kurtosis, while the homogeneity of variances was verified using Levene’s test with 5% significance. If normality and homogeneity of variances were met, one sample t-test, paired-sample t-test, two-sample t-test or one-way ANOVA with Šidák post hoc test were applied. In the case of normally distributed data in which the homogeneity of variances was violated, Welch one-way ANOVA with Games-Howell post hoc test was applied (Field, 2013). In the case of non-normally distributed variables, we performed an hyperbolic arcsine transformation, used to control for distributions with positive skewness and kurtosis that contain data with positive and negative percentage values (Tsai et al., 2017). As the transformed data verified the assumption of homogeneity of variances, two-sample t-test or one-way ANOVA with Šidák post hoc test were applied. Transformed data were only used for analytical interpretation purposes, with all figures showing the original distributions. Each test is reported with the corresponding p-value, unadjusted in case of single comparisons (analysis of joint delta power changes in each line, total NREM sleep time, NREM sleep fragmentation index and cumulative number of microarousals) or adjusted in case of multiple comparisons (analysis of delta power changes per genotype/line), with significance set to 0.05. Results are shown as the mean and SEM.

## Results

### A 2Hz component enables high NREM sleep staging precision, with 30⁰/40⁰ up-phase-targets prompting best diagnostic outcomes and ideal trigger distribution

To identify slow-waves in the ongoing EEG mice signal with the highest precision, we tested 1, 1.5, 2 and 2.5Hz frequency components in WT and TG animals (for details on the N per group, see **Supplementary Table 1**). These frequencies were coupled with a fixed phase-target of 60⁰, which previously yielded high NREM sleep staging precision in rats (Moreira et al., 2021).

We determined daily online staging precision in both genotypes in relation to the offline staging tool (Miladinović et al., 2019). In WT mice, an average of 60% epochs were confirmed offline as NREM sleep when tracking a 2Hz frequency component, whereas other frequencies showed lower precision values of 39%, 46% and 48% for 1, 1.5 and 2.5Hz, respectively (**Figure 2A**). For TG mice, we also obtained highest precision with a 2Hz component (61%), versus 43%, 57% and 60% in 1, 1.5 and 2.5Hz, respectively (**Figure 2B**). Hence, tracking higher frequency components than the one previously defined for rats (1Hz) led to high precision of online NREM sleep staging in mice.

**Figure 2.**
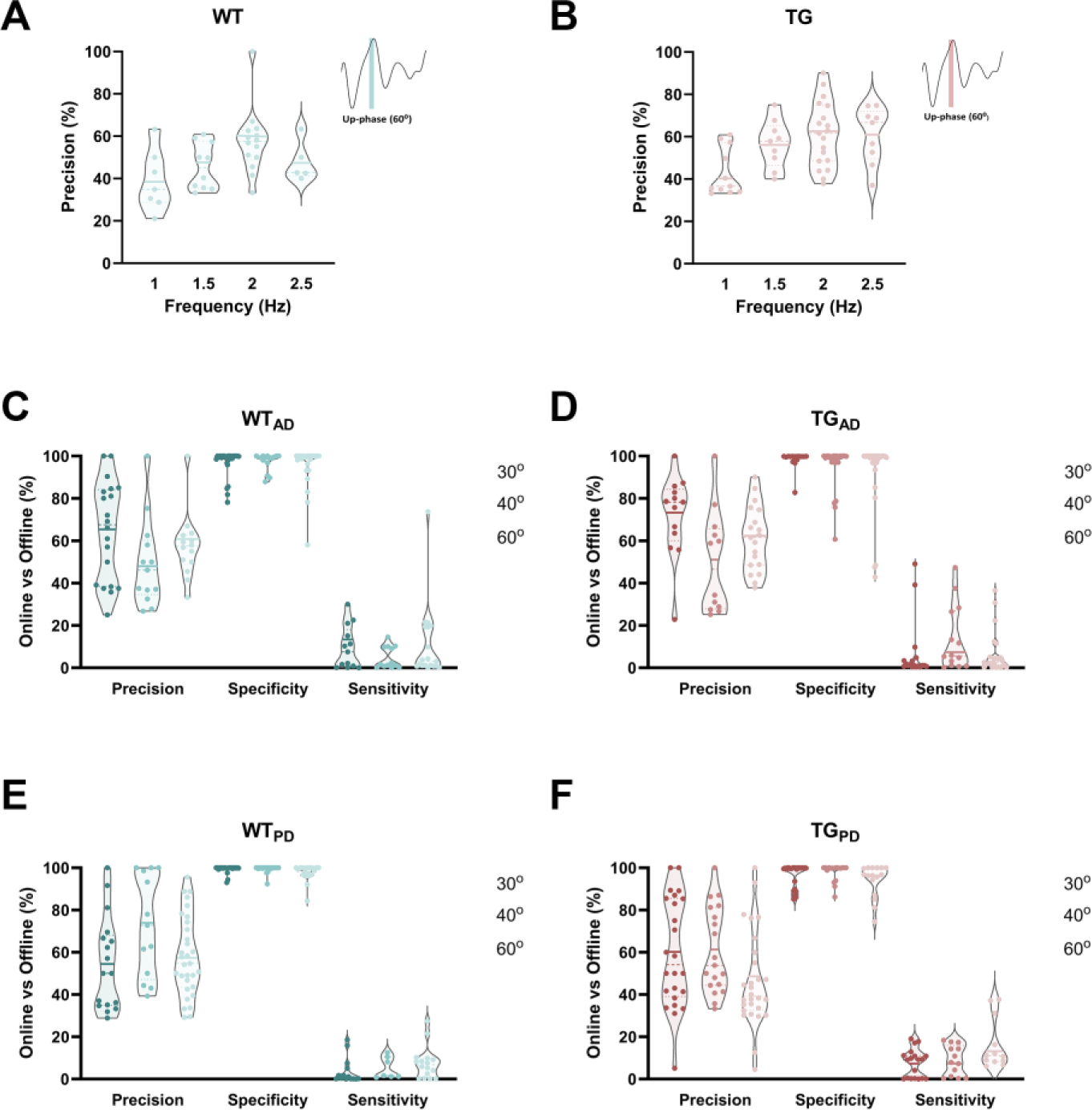
Diagnostic parameters (precision, specificity and sensitivity) for online staging performance assessment with increasing frequency components and up-phase targets. **A:** In WT mice, we obtained an average precision of 60% by tracking a 2 Hz component, in relation to 39%, 46% and 48% in 1, 1.5 and 2.5 Hz, respectively. **B:** For TG mice, we obtained an average precision of 61% by tracking a 2 Hz component, versus 43%, 57% and 60% in 1, 1.5 and 2.5 Hz, respectively. Insets depict the 60⁰ phase target used in combination with the different frequencies tested. **C:** Full set of diagnostic outcomes (precision, specificity and sensitivity) in WT_AD_ mice. **D:** Diagnostic outcomes in TG_AD_ animals. Highest precision was obtained with 30⁰ (72%), as opposed to 50% and 61% in 40⁰ and 60⁰, respectively. CLAS’ average specificity was of 98% with 30⁰ and of 95% and 94% with 40⁰ and 60⁰. Average sensitivity in TG_AD_ mice amounted to 7% with 30⁰ phase-target as opposed to 13% and 6% with 40⁰ and 60⁰. **E:** Diagnostic measures in WT_PD_ mice. **F:** Diagnostic outcomes in TG_PD_ mice. Highest precision was also obtained with 40⁰ (61%), as opposed to 60% and 46% with 30⁰ and 60⁰, respectively. Similarly, CLAS’ average specificity was of 98% with a 40⁰ and of 96% and 94% with 30⁰ and 60⁰. Average sensitivity in TG_PD_ mice amounted to 9% with 40⁰ phase-target as opposed to 8% and 16% with 30⁰ and 60⁰. Points in each violin plot represent days from all mice, dashed and dotted lines the median and quartiles, and the full blue/pink line the mean of the distributions. Original data is plotted, prior to outlier removal.

In subsequent experiments, we tracked 2Hz component to identify slow-waves in WT and TG mice.

We thereafter combined 2Hz component with three up-phase-targets of auditory stimuli delivery (30⁰, 40⁰ or 60⁰) of the ongoing slow-wave. To determine the best combination of parameters, we assessed their effect onto the full set of diagnostic outcomes (precision, specificity and sensitivity) of the online staging paradigm in WT_AD_, TG_AD_, WT_PD_ and TG_PD_ mice, independently.

In WT_AD_ mice, targeting the 30⁰ up-phase of the slow-wave led to highest precision of online NREM sleep detection (65%) as opposed to 49% and 60% for 40⁰ and 60⁰, respectively (**Figure 2C**). In these mice, mCLAS’ average specificity was 97% with 30⁰ phase-target and of 97% and 96% with 40⁰ and 60⁰, respectively. Average sensitivity in WT_AD_ mice was highest with 30⁰ phase-target (12%) as opposed to 5% and 10% with 40⁰ and 60⁰, respectively (**Figure 2C**). We obtained similar average values for precision, specificity and sensitivity in TG_AD_ mice (**Figure 2D**).

Contrastingly, in WT_PD_ mice we obtained highest precision with 40⁰ phase-target (73%) as opposed to 56% and 58% with 30⁰ and 60⁰, respectively (**Figure 2E**). In these mice, mCLAS’ average specificity was of 99% with 40⁰ and of 99% and 98% with 30⁰ and 60⁰, respectively. Average sensitivity in WT_PD_ mice amounted to 6% with 40⁰ phase-target as opposed to 5% and 8% with 30⁰ and 60⁰, respectively (**Figure 2E**). We obtained similar average values for all diagnostic outcomes in TG_PD_ mice (**Figure 2F**).

Upon assessing that a parameter combination of 2Hz/30⁰ (WT_AD_ and TG_AD_) or 2Hz/40⁰ (WT_PD_ and TG_PD_) yielded the best set of diagnostic outcomes within the lines, we calculated the percentage of total triggers delivered and the daily number of triggers in NREM sleep and other vigilance stages per animal, with these parameter combinations. In WT_AD_ mice, an average of 58% of total triggers per animal per day were delivered during NREM sleep, similarly to 60% observed in TG_AD_ mice (**Figure 3A**). We also obtained an average of 3382±519 daily triggers in NREM sleep for WT_AD_ mice, which contrasted with 2161±397 daily triggers in NREM sleep for TG_AD_ mice (**Figure 3B**). In WT_PD_ animals, 61% of triggers were delivered during NREM, whereas 51% were delivered in TG_PD_ (**Figure 3C**). An average of 2771±555 triggers were delivered daily in WT_PD_ whereas 4633±606 triggers were delivered in TG_PD_ mice (**Figure 3D**).

**Figure 3.**
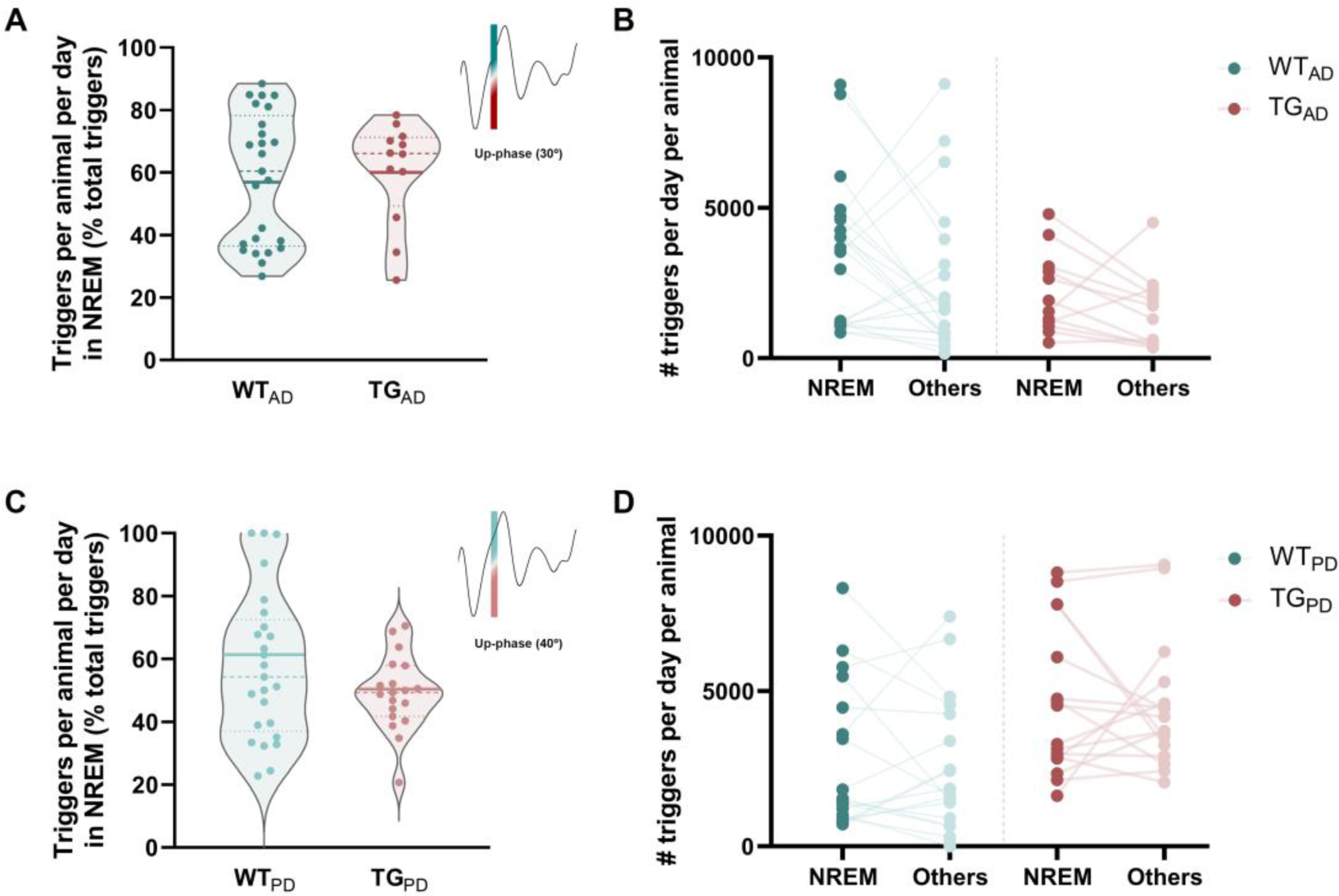
Percentage of daily triggers delivered during NREM sleep per animal, and daily number of triggers in NREM versus other vigilance stages in WT_AD_, WT_PD_, TG_AD_ and TG_PD_ mice. **A:** Triggers per animal per day as percentage of total triggers during NREM sleep in WT_AD_ and TG_AD_ mice. **B:** Average daily triggers in NREM sleep for WT_AD_ and TG_AD_ mice. **C:** Triggers per animal per day as percentage of total triggers during NREM sleep in WT_PD_ and TG_PD_ mice. **D:** Average daily triggers in NREM sleep for WT_PD_ and TG_PD_ mice. Insets in left panels depict the phase target analyzed within each measure. Points in each violin plot represent days from all animals, full blue/red line the median and dotted lines the quartiles.

To assess the ability of the mCLAS paradigm to deliver stimuli across the up-phase-targets tested combined with a 2Hz component, we averaged the instantaneous phase for all triggers delivered in NREM sleep, appraising trigger distribution along the slow-wave for each phase-target. In WT_AD_ and TG_AD_ mice, the majority of triggers targeted at 30⁰ and 40⁰ up-phases of the slow-wave fell within its rising phase (**Figure 4A,B**). The highest percentage of triggers is distributed along up-phases immediately before the positive peak, in the 50⁰-90⁰ range. In contrast, with 60⁰ phase-percentage distributions show heavy right tales (>90⁰) in these mice (**Figure 4C**). We observed similar trigger distribution patterns for WT_PD_ and TG_PD_ mice (**Figure 4D-F**).

**Figure 4.**
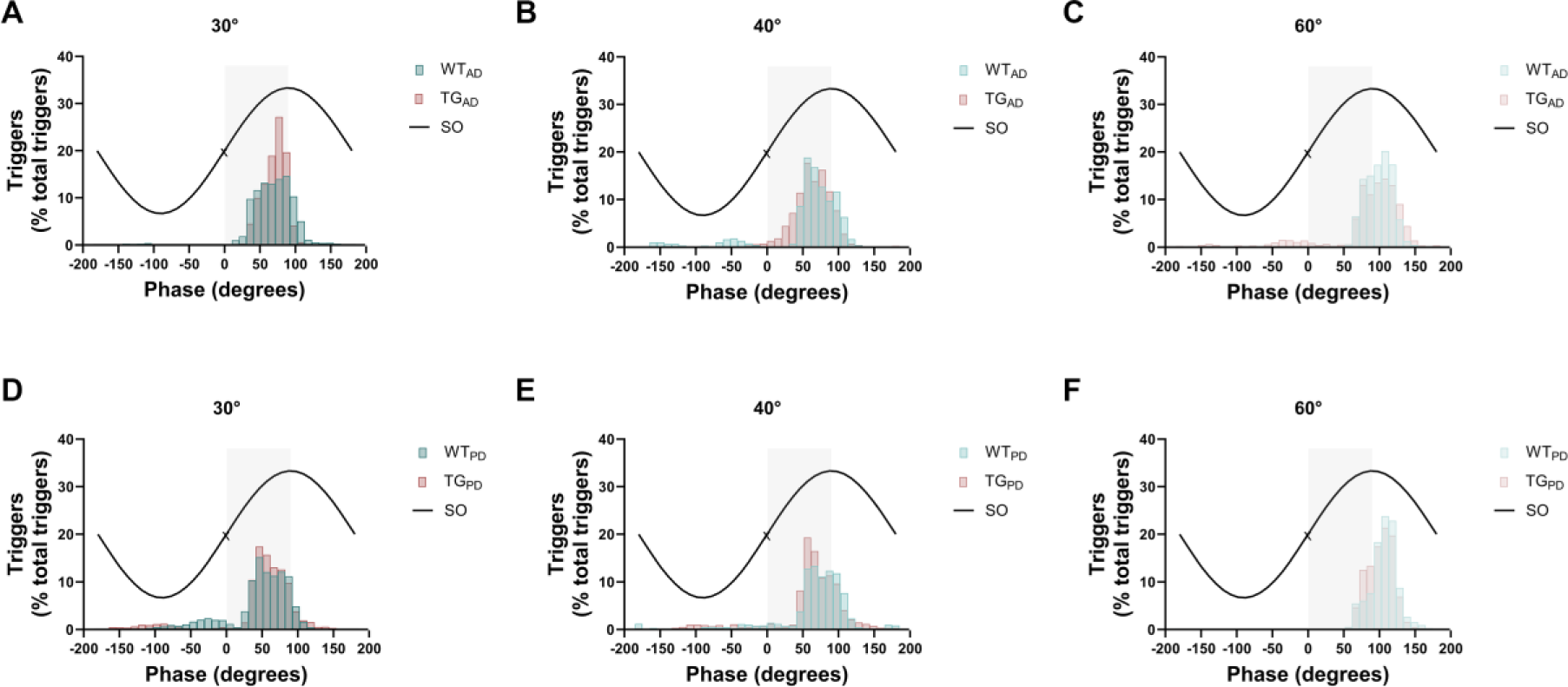
Distribution of triggers for each up-phase target and mouse model. **A-C:** Trigger distributions in the AD model (WT_AD_ and TG_AD_) per up-phase target. **D-F:** Trigger distributions in the PD model (WT_PD_ and TG_PD_) per up-phase target. Histograms are depicted with bins of 10°. Counts were normalized to the total number of counts in each animal. SO – slow oscillation; \ - SO negative-to-positive zero crossing; Grey background – rising phase (0⁰ - 90⁰).

### Phase-targeted mCLAS evokes genotype independent but strain specific delta power changes in mice

After establishing optimal stimulation parameters for each strain, we aimed to verify mCLAS’ ability to enhance sleep depth in both neurodegeneration murine models. We calculated power changes in the delta range as percentage of change from baseline per animal and per day of stimulation, for each phase-target combined with a 2Hz component.

In WT_AD_ mice, we observed a significant delta power increase after stimulation with 30⁰ phase-target compared with the MOCK group (29±8%, p<0.001, one-way ANOVA, Šidák correction; **Figure 5A**). Other phase-targets assessed in this genotype led to decreases in delta power, which were significantly different from the increase with 30⁰ (40⁰: -12±4%, p<0.001; 60⁰: -9±4%, p<0.001). On the other hand, delta power in WT_PD_ was significantly increased upon stimulation with a phase-target of 40⁰ (35±21%, p=0.053) compared with MOCK. mCLAS-evoked delta power decreases with 30⁰ (-6±6%, p<0.01) and 60⁰ (-2±8%, p=0.057) differed from the increase with 40⁰ (**Figure 5A**).

**Figure 5.**
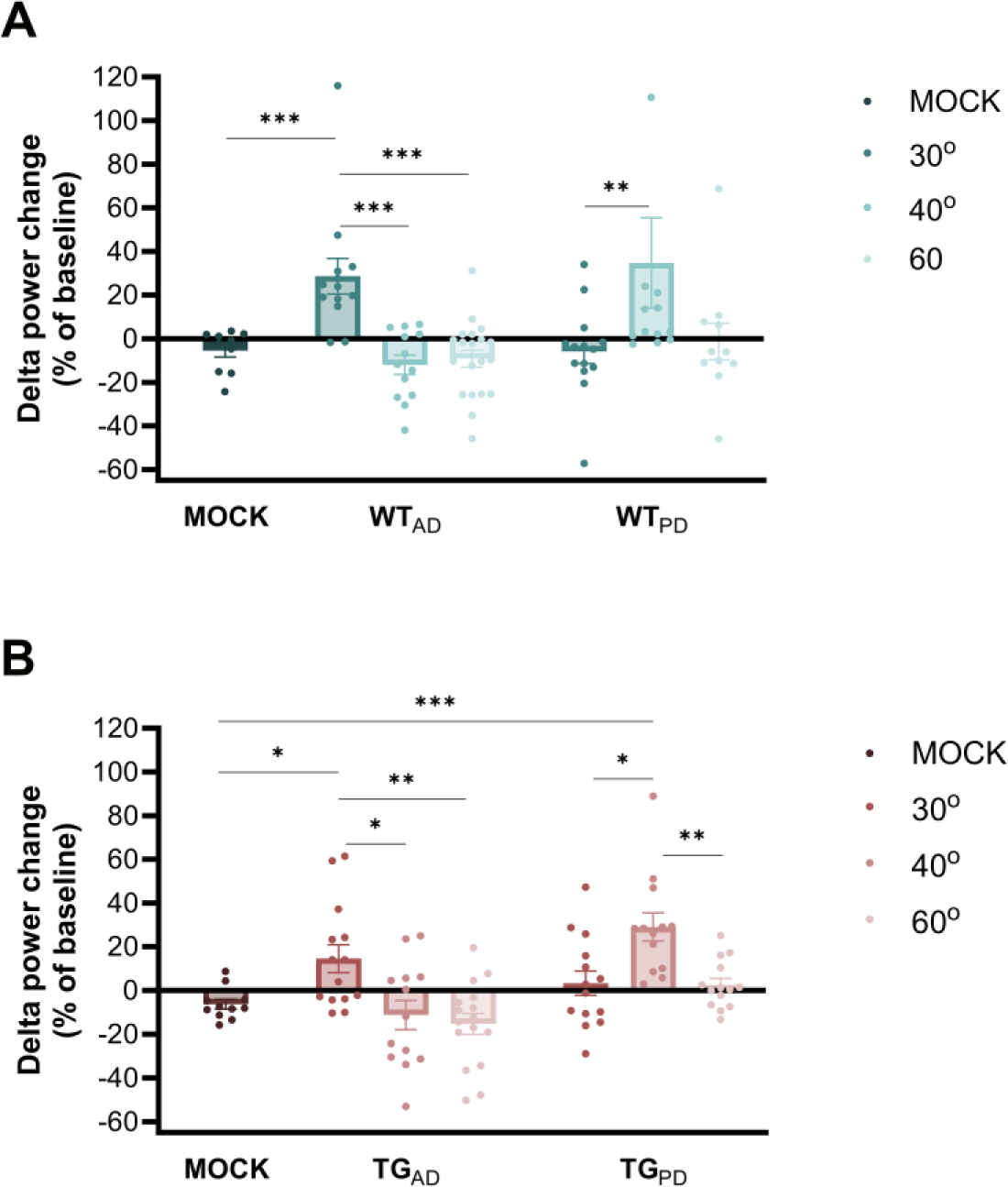
Delta power changes evoked by 30⁰, 40⁰ and 60⁰ up-phase targets in MOCK, WT_AD_, WT_PD_, TG_AD_ and TG_PD_ mice. **A:** Delta power changes in WT_AD_ and WT_PD_ mice. **B:** Delta power changes in TG_AD_ mice and TG_PD_ mice. **C:** Superimposed distributions of delta power changes of MOCK, WT_AD_ with TG_AD_, and WT_PD_ with TG_PD_ mice confirm similar patterns of delta power changes within each joint distribution across up-phase targets. Each scatter dot represents 24h of delta power change as percentage of baseline of each animal, each bar the mean of such changes, and the whiskers the SEM. ***p<0.001, **p<0.01, *p<0.05.

For TG_AD_ mice, we observed a significant delta power increase upon 30⁰ stimulation (15±6%, p<0.05, one-way ANOVA, Games-Howell correction; **Figure 5B**) in comparison with MOCK. Again, 40⁰ and 60⁰ targets led to significant delta power decreases, which significantly different from the increase with 30⁰ (40⁰: -11±7%, p<0.05; 60⁰: -15±5%, p<0.01). Conversely, in TG_PD_ animals delta power was increased after stimulation with all up-phase-targets when compared to MOCK, but only significantly with 40⁰ (31±6%, p<0.001). Other up-phase-targets yielded modest delta power increases, which significantly differed from that elicited by 40⁰ (30⁰: 3±6%, p<0.05; 60⁰: 1±3%, p<0.01; **Figure 5B**).

Upon observing similarities in the direction of mCLAS-induced delta power changes between genotypes of the same model, we superimposed the distribution of delta power change values of MOCK, and up-stimulated WT_AD_ with TG_AD_, and WT_PD_ with TG_PD_ mice. We confirmed largely overlapping patterns of delta power change values within each joint distribution across up-phase-targets **(Supplementary Figure 1A)** with no significant within differences, suggesting no genotype-dependent effect.

To infer on the possible relationship between genetic strains and target-specific mCLAS-evoked delta power changes, we pooled delta power change values for each strain, obtaining data point distributions for AD (B6;SJL) and PD (B6.Cg) lines. We compared these joint distributions within each up-phase-target and observed that the distributions for each strain significantly differed within all targets (**Supplementary Figure 1B**). In B6;SJL mice, we found that the average 22±5% delta power increase with 30⁰ phase-target significantly differed from -2±5% decrease in B6.Cg (p<0.01, two-sample t-test), whereas the -12±4% decrease obtained with 40⁰ in B6;SJL significantly differed from a 32±12% increase in B6.Cg (p<0.001). Finally, in B6;SJL mice, we detected a -12±3% delta power decrease with 60⁰, which was significantly different from the -1±3% decrease observed in B6.Cg mice (p<0.05).

### mCLAS elicits a decrease in pathologically-heightened NREM sleep fragmentation and microarousal events in transgenic AD mice

After determining that short-term application of mCLAS leads to significant delta power increases in neurodegeneration models, we aimed at exploring its potential effects onto disease-triggered sleep alterations.

In terms of macrostructural sleep architecture, we assessed total time spent in NREM sleep in minutes and as percentage of change from baseline, for each animal. In TG_AD_ mice, we observed a decrease in total time spent in NREM sleep at baseline, in relation to WT_AD_ animals (**Figure 6A**). This initial impairment was not significantly counteracted by mCLAS. Similarly, in WT_AD_ mice the stimulation paradigm exerted no modulatory effect on total time spent in NREM sleep. In both WT_PD_ and TG_PD_ mice, total NREM sleep time remained fairly constant throughout the stimulation days, not changing significantly from non-altered baseline values (**Figure 6B**). Moreover, we calculated NREM sleep fragmentation index as a percentage of change from baseline, for each animal. At baseline, NREM sleep fragmentation was higher in TG_AD_ mice in relation to WT_AD_ mice (**Figure 6C**). This disease-specific impairment was counteracted by mCLAS, which elicited a significant 14% decrease in NREM sleep fragmentation of TG_AD_ animals when compared to baseline values (p<0.01, one-sample t-test). NREM sleep fragmentation index remained unchanged in WT_AD_ mice. Contrastingly, we observed no initial impairment in NREM sleep fragmentation of TG_PD_ mice, in relation to their WT_PD_ littermates (**Figure 6D**) and, concomitantly, mCLAS exerted no significant modulatory effect in NREM sleep fragmentation in this strain.

**Figure 6.**
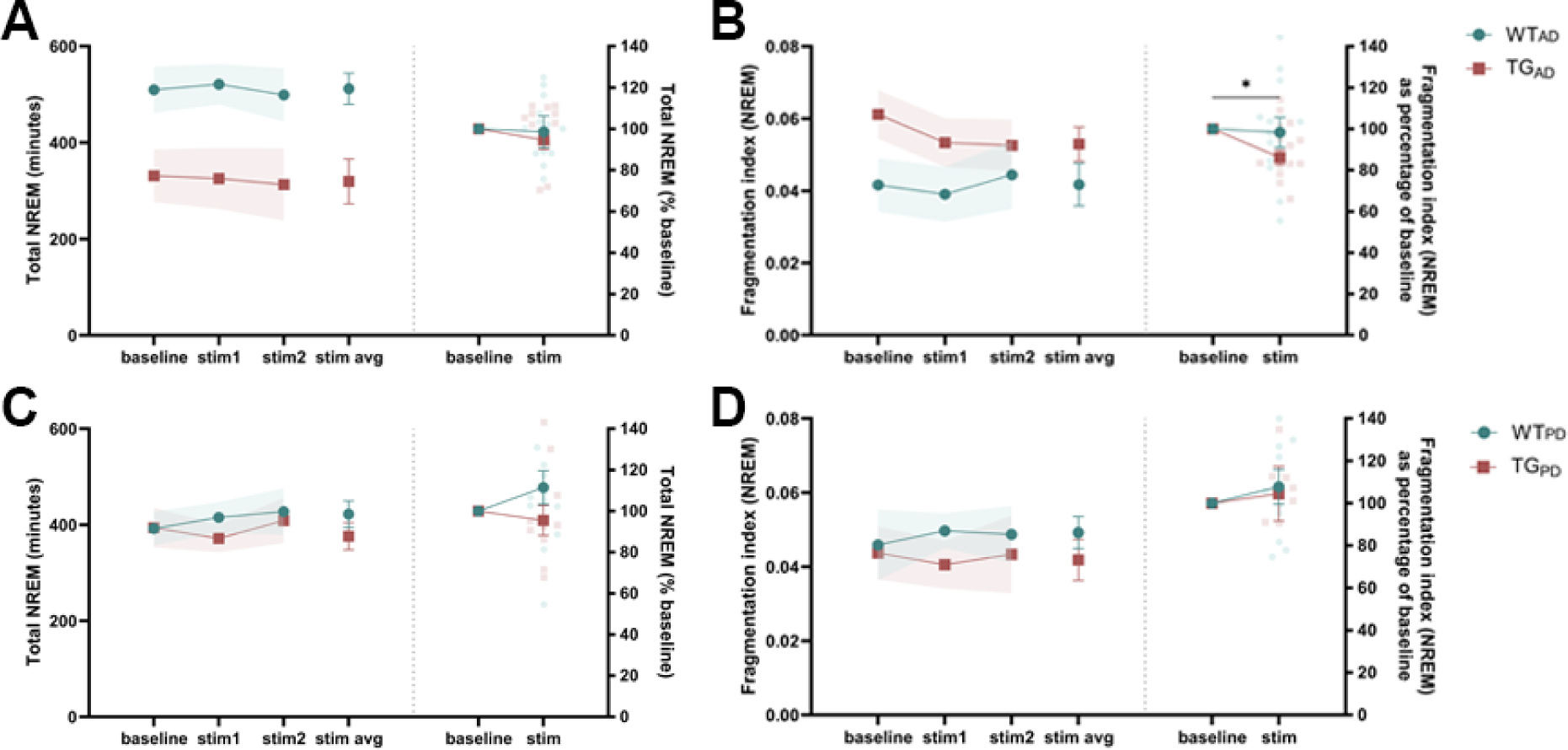
Total NREM sleep duration and fragmentation index in the AD and PD model. **A:** Total time spent in NREM sleep (in minutes) of WT_AD_ and TG_AD_ mice. In both genotypes, no changes were observed between baseline and mCLAS (30⁰ up-phase target). **B:** Total time spent in NREM sleep (in minutes) of WT_PD_ and TG_PD_ mice. In both genotypes, no changes were observed between baseline and mCLAS (40⁰ up-phase target). **C:** mCLAS significantly decreases a pathologically heightened NREM sleep fragmentation index in TG_AD_ mice but not in WT_AD_ animals. **D:** Fragmentation index in both WT_PD_ and TG_PD_ mice remains unchanged between baseline recordings and mCLAS. Shaded areas correspond to the SEM. **p<0.01.

Finally, as a metric of sleep depth and maintenance (dos Santos Lima et al., 2019; Gradwohl et al., 2017; Sforza et al., 2004), we assessed microarousal distributions across 24h and cumulatively in 24h, light and dark periods, in WT_AD_, TG_AD_, WT_PD_ and TG_PD_ mice. In WT_AD_ mice, mCLAS elicited a non-significant decrease in microarousals during the dark period in relation to baseline values (**Figure 7A**- **B**). Interestingly, in TG_AD_ animals, mCLAS lead to a significant decrease in the number of microarousals across both light and dark periods, when compared with pathologically-heightened baseline values (p<0.05, paired sample t-test; **Figure 7C-D**). Contrastingly, mCLAS did not significantly alter daily microarousal dynamics in WT_PD_ and TG_PD_ mice, with both distributions and cumulative number of microarousals remaining constant throughout the stimulation procedure, maintaining similar non-pathological patterns observed at baseline (**Figure 7E-H**).

**Figure 7.**
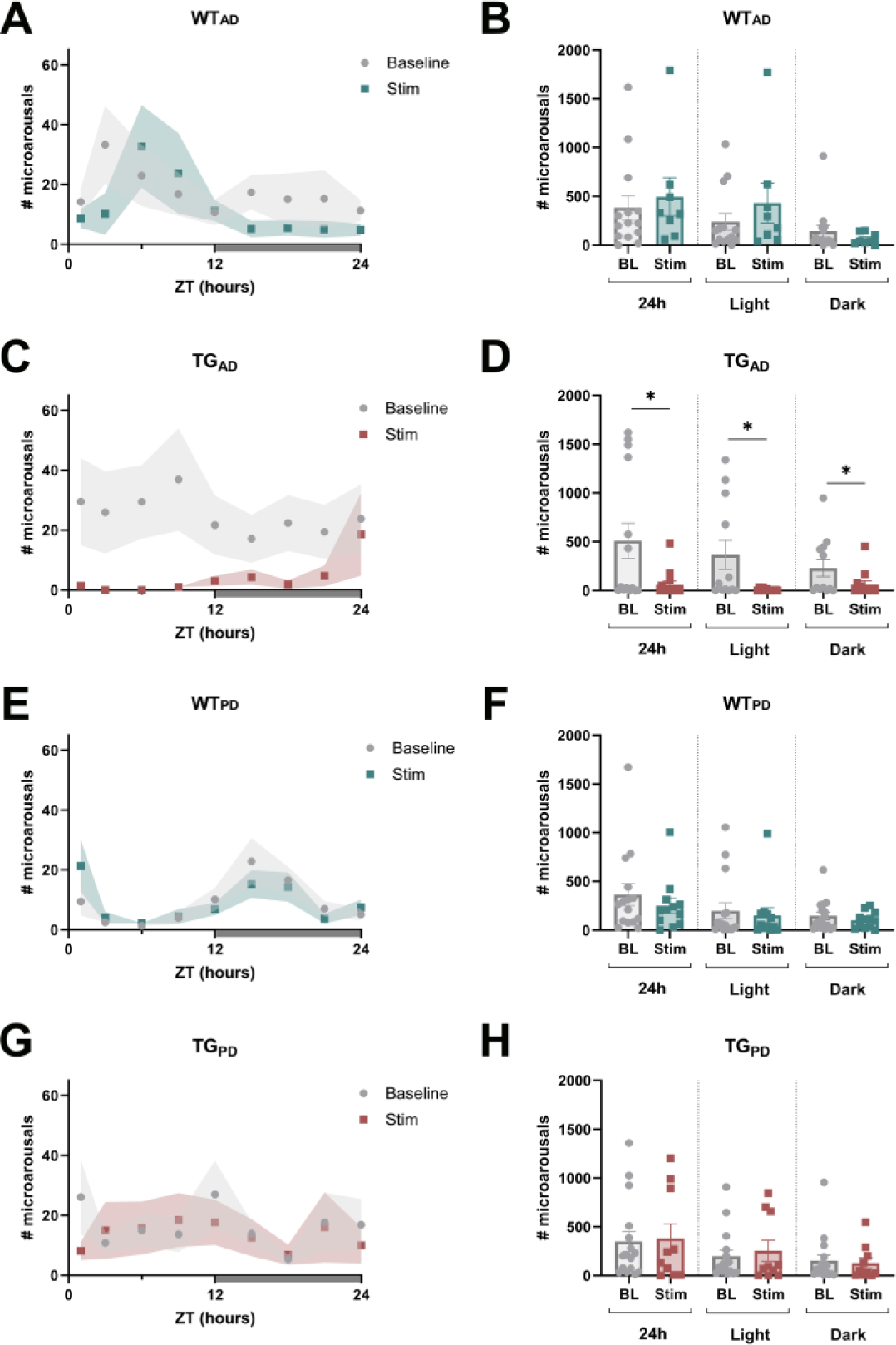
Microarousal dynamics across 24h and cumulative number of microarousals per 24h, light and dark periods. **A**: In WT_AD_ mice, mCLAS (30⁰ up-phase target) leads to a decrease in microarousal dynamics during the dark period in comparison with baseline recordings, which is not significant (**B**). **C**: In TG_AD_ mice, mCLAS elicits a decrease in microarousal dynamics in both light and dark periods in comparison with baseline dynamics. This decrease is significant across 24h, light and dark periods (**D**). **E, F, G** & **H**: In WT_PD_ and TG_PD_ mice, mCLAS (40⁰ up-phase target) did not significantly modulate microarousal dynamics across 24h nor the cumulative number of microarousals across 24h, light and dark periods. Shaded areas correspond to the SEM. In the 24h dynamics plots, the grey shaded bar corresponds to the dark period. In the bar plots, each bar represents the mean cumulative number of microarousals, and the whiskers the SEM. *p<0.05.

## Discussion

In this study, we established up-phase-targeted mCLAS in mouse models of neurodegeneration by developing a paradigm applicable to wildtype and transgenic animals. To achieve this, we tracked four frequency components in the EEG signal and targeted three up-phases within the slow-wave. We determined that 2Hz/30⁰ or 2Hz/40⁰ revealed optimal diagnostic outcomes and trigger distributions along the up-phase of the slow-wave in each line, which translated into delta power increases that were not genotype-dependent but strain-specific. Importantly, these short-term mCLAS-evoked delta power increases prompted alleviation of neurodegeneration-associated sleep alterations in AD mice.

### CLAS is valid and applicable in mouse models of neurodegeneration

We successfully implemented a novel paradigm of mCLAS in healthy and diseased mice within the context of neurodegeneration. We have previously developed this technique in healthy and traumatic brain injury rats (Moreira et al., 2021, 2022), using similar CLAS’ parameters proposed in humans (Fattinger et al., 2017; Ngo et al., 2013), as slow-wave characteristics appear to be fairly similar between those two species (Mölle et al., 2009). In contrast, slow-waves in mice have shorter wavelengths and peak at higher frequencies within the delta range (Panagiotou et al., 2017). Accordingly, we obtained on average 61%-73% precision by adjusting the EEG tracked frequency to 2 fold that of rats and targeting earlier slow-wave phases. These precision values were the highest among the different parameters tested and are comparable with the staging precision previously obtained in rats and humans (Fattinger et al., 2017; Moreira et al., 2021; Papalambros et al., 2017). Specificity and sensitivity were also fairly comparable with previous reports, illustrating the high rejection rate of events without NREM characteristics, and how strictly the online staging paradigm identifies sustained NREM sleep. Moreover, the 2Hz/30⁰ and 2Hz/40⁰ parameter combinations revealed trigger distributions that overlap with the rising phase of the slow-wave close to its positive peak, mimicking the distribution obtained with the 1Hz/60⁰ combination in rats (Moreira et al., 2021). This up-phase overlapping is reflected in a steady increase in delta power upon stimulation, ranging 15 and 35% when compared to a MOCK condition. This increase is between 2-5 fold higher than the increase previously observed in healthy rats upon up-phase stimulation (∼7%), stressing the importance of tailoring CLAS’ parameters to maximize its effect.

Literature on human CLAS also advocates for precise phase-targeting when aiming at boosting SWA during deep sleep (Krugliakova et al., 2022; Ngo et al., 2013), although it has often failed to demonstrate an overall inter-night effect of stimulation on delta power (baseline nights vs. stimulation nights), recurring to comparison of same-night ON-OFF windows (Krugliakova et al., 2022; Papalambros et al., 2017, 2019). Our results using a continuous stimulation paradigm suggest that the effect of mCLAS is due to a general SWA enhancement and not mediated by qualitative transient reorganization of SWA, as proposed elsewhere (Papalambros et al., 2017). In turn, homeostatic compensatory mechanisms might be at play (Kim et al., 2020), meaning our mCLAS paradigm may lead to an increased homeostatic decline mediated by an increase in SWA during deep sleep (Krugliakova et al., 2022).

The percentage of daily delivered triggers in NREM sleep is also comparable to previous results in rats (Moreira et al., 2021), amounting for an average 51-61% of total triggers. Auditory stimuli would ideally be delivered only during sustained NREM, but rhythmic motion behaviors such as grooming, feeding, and drinking may have majorly contributed to non-NREM epochs being staged online as NREM sleep. Interestingly, we observed a lower number of auditory triggers delivered during NREM in TG_AD_ mice in relation to WT_AD_. This observation aligns with the previously reported reductions in NREM sleep during the dark period and number of slow oscillations in 7-month-old TG_AD_ mice of the Tg2576 line (Kollarik et al., 2022). Contrastingly, we observed a greater number of triggers during NREM sleep in TG_PD_ mice in relation to WT_PD_. To our knowledge, only one study has quantitively reported on sleep characteristics in the A53T line, describing a decrease in total NREM sleep time in 7-month-old TG_PD_ animals in comparison to WT_PD_, which co-occurred with increased delta power in the mutants and hyperexcitability with epileptiform events following slow-waves (Peters et al., 2020). These results agree with our observations, as increased delta power coupled to high amplitude events might have led to increased number of triggers during NREM sleep in TG_PD_ mice.

Potentially altered slow-wave characteristics associated to neurodegeneration could have affected mCLAS’ ability to target and stimulate SWA. Such was not the case observed in our paradigm, likely indicating that mCLAS is capable of bypassing disease-associated slow-wave alterations and still successfully track and stimulate them.

### mCLAS-evoked phase-specific changes depend on the genetic background across transgenic strains

Our mCLAS paradigm successfully induced up-phase-specific delta power changes independently of genotype, but in turn determined by the genetic strain of the transgenic neurodegeneration model.

The two transgenic murine models assessed in this study (AD: Tg2576 line; PD: M83 line) were bred in different backgrounds (AD: B6;SJL; PD: C3H/HeJ x C57BL6/J), resulting in distinct murine strains (Giasson et al., 2002; Hsiao et al., 1996). Although literature directly comparing these specific mouse lines is lacking, several studies have reported on strain-dependent auditory processing differences between the genetic backgrounds that compose the two models. Auditory brainstem responses in SJL and C3H/HeJ strains were reported to have different inter-peak latencies -being those longer in C3H/HeJ -, implying distinct synaptic delays and neural conduction velocities within the brainstem pathways (Zhou et al., 2006). Moreover, recording of auditory-evoked potentials revealed a response to novel and standard tones in the C3H/HeJ strain that was absent in C57BL/6J mice, suggesting that these strains differ in their novelty responses and ability to neuronally detect changes in the auditory environment (Siegel et al., 2003). Upon auditory stimulation in mice, our results also suggest that genetic strain is determinant for informing optimal parameter choice when aiming at modulating SWA. Overall, our results hint that genetic background could have a significant role in determining the ideal phase-target necessary to observe a mCLAS-evoked delta power increase, perhaps due to affected velocities of neural processing, slow-wave characteristics and/or other relevant variables. To our knowledge, this is the first account of genetic dependency in the context of phase-targeted CLAS, with potential major implications for CLAS human studies and future clinical implementations.

### mCLAS rescues altered sleep consolidation in AD mice

AD mice present heightened NREM sleep fragmentation accompanied by a decrease in NREM sleep duration compared to WT animals (Kollarik et al., 2022). Increased sleep fragmentation is a distinctive feature of neurodegenerative diseases, being linked to accelerated microglia and astrocyte aging and activation, cognitive impairment in patients, increasing amyloid-beta deposition, tau phosphorylation and neuroinflammation, and decreasing AQP4 expression in older AD animals (Cai et al., 2019; Kaneshwaran et al., 2019; Vasciaveo et al., 2023). Consequently, sleep alterations are clinical hallmarks that appear as crucial risk factors in neurodegeneration, and should be explored as therapeutic targets to control disease progression.

A striking effect of our short-term mCLAS paradigm in TG_AD_ animals was the prompting of a 14% decrease in NREM sleep fragmentation index, and a steep decrease in microarousal events in relation to pathologically increased baseline values. These results indicate that mCLAS corrected a sleep impairment in TG_AD_ mice, perhaps via impeded NREM-REM or NREM-WAKE transitions and overall increased coherence (dos Santos Lima et al., 2019; Gradwohl et al., 2017). On the other hand, microarousals represent a transient cortical activation, reflecting the brain’s response to stimuli (Sforza et al., 2004). Therefore, as a consequence of a higher sleep pressure due to increased sleep fragmentation in TG_AD_ mice, mCLAS-mediated facilitation of sleep maintenance and consolidation in these animals may have prompt a reduction of cortical responsiveness to arousing stimuli, thus resulting in lower microarousal levels. This implicates that mCLAS might lead to faster sleep dissipation via higher neuronal population synchrony, a speculation that should be verified in future studies by analyzing pre- and post-stimulation slow oscillation characteristics in more detail (Bernardi et al., 2018; Fattinger et al., 2014; Vyazovskiy et al., 2009). Moreover, it was previously hypothesized that the decrease in cortical microarousals after manipulation of homeostatic sleep pressure – perhaps elicited via mCLAS in our paradigm – would be greater than the decrease in subcortical arousals (as K-bursts and D-bursts) (Sforza et al., 2004). Thus, even though we did not directly assess subcortical arousals in this study, our results might hint that mCLAS modulates the number of microarousal events through a direct effect on cortical excitability, with slighter influence on brainstem sleep generators.

Finally, TG_PD_ mice displayed no baseline impairments in NREM sleep fragmentation and duration compared to their WT controls, partially confirming earlier evidence on the lack of abnormal sleep fragmentation in PD mice (Morawska et al., 2021). Perhaps as a consequence of these absent alterations, NREM sleep fragmentation and microarousal events remained unchanged throughout mCLAS days, suggesting that this intervention selectively modulates altered sleep stability, allowing for a highly specific alleviation of neurodegeneration-associated sleep phenotypes.

### Limitations

A methodological limitation is the mixed cohort included in the MOCK groups. Due to a low number of animals with complete datasets from MOCK in each genotype, we recurred to pooling WT_AD_ with WT_PD_ mice within the WT MOCK group, and TG_AD_ with TG_PD_ mice within the TG MOCK group. However, delta power changes and standard errors within each MOCK group appeared negligible evidencing no changes compared to baseline, suggesting no effect of mCLAS paradigm (sound insulation housing condition, etc) in these non-stimulated animals, independently of their strain. We have not performed sex-based analysis in this study, but we believe that the female-male percentage included (60-40%) balances possible sex differences. Data loss in several mice also prevented repeated measures analysis. This caveat was mainly due to software crashes, lost EEG implants, disconnections or poor signal quality associated with long-term recordings. Moreover, due to our 4s-epoch sleep scoring method, we might have not been able to observe an increase in NREM sleep duration in TG_AD_ mice upon a decrease in the number of microarousals, as these events were two small in duration (< 2 seconds) to influence scored epochs. Lastly, we did not implement adaptive feedback methods to estimate ongoing phase-targets in our experiments (Ong et al., 2016; Papalambros et al., 2017, 2019; Santostasi et al., 2016), which could potentially improve applicability of long-term mCLAS during deep sleep.

## Conclusions

In summary, we show that phase-targeted mCLAS can be successfully implemented in healthy and diseased mice in a precise manner. Some key pathological sleep traits are rescued by mCLAS, prompting the acute alleviation of neurodegeneration-associated sleep phenotypes by potentiating sleep maintenance. This novel noninvasive and nonpharmacological tool paves the way for experiments unraveling the mechanisms underlying mCLAS’ modulation on SWA, its potential long-term effect on brain disease, and the role of SWA in toxic protein accumulation processes in neurodegeneration. Hence, mCLAS may become the future cornerstone of SWA-based therapies for neurodegenerative diseases.

## Data availability

All data generated or analyzed during this study are included in the manuscript and supporting files.

## Competing interests

The authors declare no conflict of interests.

## Acknowledgments and funding statement

We would like to thank Ami Beuret, Mark Hanus, Myles Billard and Max Hunger for their valuable support. This project was funded by Parkinson Schweiz (ID, DN), the Dementia Research Switzerland - Synapsis Foundation via an earmarked donation of the Armin and Jeannine Kurz Stiftung (DN), the Neuroscience Center Zurich via a donation of Rahn & Bodmer banquiers (DN), the Hurka Foundation (DN) and the Swiss National Science Foundation (CRB). The funding sources had no involvement in study design, collection, analysis, interpretation of the data nor in writing or deciding to submit this article for publication.

## Author contributions

**Inês Dias**: Conceptualization, Methodology, Software, Validation, Formal Analysis, Investigation, Data Curation, Visualization, Funding acquisition, Writing – Original Draft, Writing – Review & Editing; **Sedef Kollarik**: Formal Analysis, Investigation, Writing - Review & Editing; **Michelle Siegel**: Investigation, Writing – Review & Editing; **Christian R. Bauman**: Funding acquisition, Writing – Review & Editing; **Carlos G. Moreira**: Software, Writing – Review & Editing; **Daniela Noain**: Conceptualization, Resources, Supervision, Project administration, Funding acquisition, Writing – Original Draft, Writing – Review & Editing.

## Supplementary tables and figures

**Table 1.** Number of mice recorded per frequency, phase and genotype. **Top:** all frequencies were tested in combination with a phase target of 60⁰, a phase target previously successful in rats; Bottom: all phase targets tested were combined with a tracked 2 Hz frequency component. Number of females per group are provided in brackets.

|  | Frequency (Hz) |  |  |  |
| --- | --- | --- | --- | --- |
|  | 1 | 1.5 | 2 | 2.5 |
| **WT** | 5 (3) | 6 (2) | 10 (6) | 5 (3) |
| **TG** | 5 (3) | 7 (4) | 10 (6) | 5 (3) |

|  | Phase (°) |  |  |  |
| --- | --- | --- | --- | --- |
|  | MOCK | 30 | 40 | 60 |
| **WT<sub>AD</sub>** | 4 (1) | 7 (5) | 7 (5) | 10 (6) |
| **WT<sub>PD</sub>** | 3 (2) | 7 (5) | 7 (5) | 6 (4) |
| **TG<sub>AD</sub>** | 4 (2) | 7 (4) | 8 (4) | 10 (6) |
| **TG<sub>PD</sub>** | 4 (3) | 7 (6) | 7 (6) | 7 (6) |

**Supplementary Figure 1.**
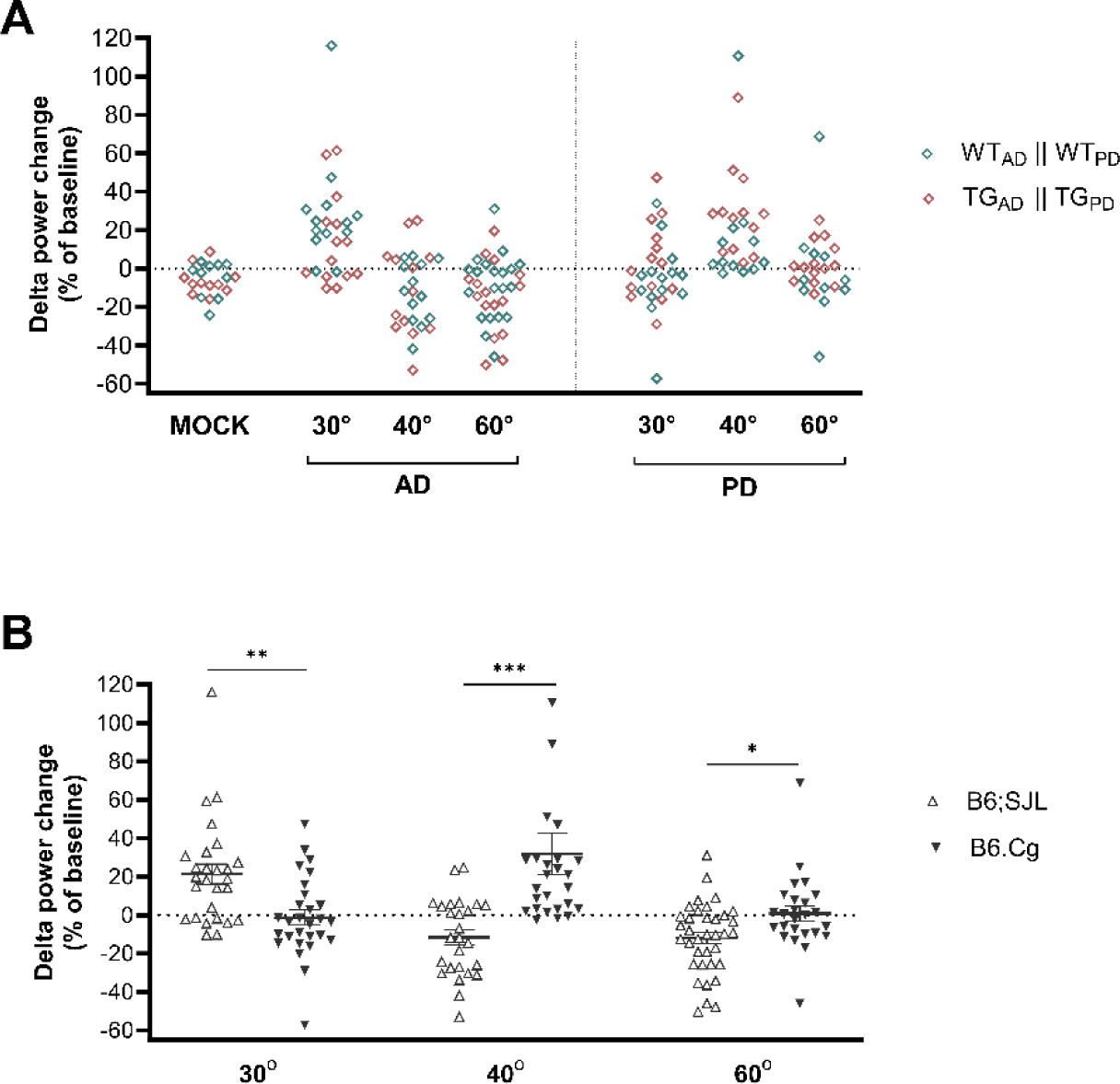
Delta power changes for B6;SJL (WT_AD_ + TG_AD_) and B6.Cg mice (WT_PD_ + TG_PD_). **A:** Superimposed distributions of delta power changes of MOCK, WT_AD_ with TG_AD_, and WT_PD_ with TG_PD_ mice confirm similar patterns of delta power changes within each joint distribution across up-phase targets. **B:** Distributions of CLAS-evoked delta power changes within all u-phase targets for each mouse strain. Scatter dots represent delta power change per 24h as percentage of baseline of each animal. Solid black horizontal line correspond to the mean per group, and the whiskers the SEM. ***p<0.001, **p<0.01, *p<0.05.

## Notes

### Competing Interest Statement

The authors have declared no competing interest.

